# Dedifferentiated early postnatal lung myofibroblasts redifferentiate in adult disease

**DOI:** 10.1101/2023.10.04.560924

**Authors:** Rachana R. Chandran, Taylor Adams, Inamul Kabir, Eunate Gallardo, Naftali Kaminski, Brigitte Gomperts, Daniel M. Greif

**Affiliations:** Yale Cardiovascular Research Center, Section of Cardiovascular Medicine, Department of Medicine, Yale University School of Medicine, New Haven, CT, USA; Department of Genetics, Yale University School of Medicine, New Haven, CT, USA; Section of Pulmonary, Critical Care and Sleep Medicine, Department of Internal Medicine, Yale University School of Medicine, New Haven, CT, USA; Children’s Discovery and Innovation Institute, Mattel Children’s Hospital, Department of Pediatrics, UCLA, Los Angeles, CA, USA; Division of Pulmonary and Critical Care Medicine, David Geffen School of Medicine, UCLA, Los Angeles, CA, USA; Jonsson Comprehensive Cancer Center, UCLA, Los Angeles, CA, USA; Eli and Edythe Broad Stem Cell Research Center, UCLA, Los Angeles, CA, USA; Molecular Biology Institute, UCLA, Los Angeles, CA, USA

## Abstract

Alveolarization ensures sufficient lung surface area for gas exchange, and during bulk alveolarization in mice (postnatal day [P] 4.5-14.5), alpha-smooth muscle actin (SMA)^+^ myofibroblasts accumulate, secrete elastin, and lay down alveolar septae. Herein, we delineate the dynamics of the lineage of early postnatal SMA^+^ myofibroblasts during and after bulk alveolarization and in response to lung injury. SMA^+^ lung myofibroblasts first appear at ∼P2.5 and proliferate robustly. Lineage tracing shows that, at P14.5 and over the next few days, the vast majority of SMA^+^ myofibroblasts downregulate smooth muscle cell markers and undergo apoptosis. Of note, ∼8% of these dedifferentiated cells and another ∼1% of SMA^+^ myofibroblasts persist to adulthood. Single cell RNA sequencing analysis of the persistent SMA^-^ cells and SMA^+^ myofibroblasts in the adult lung reveals distinct gene expression profiles. For instance, dedifferentiated SMA^-^ cells exhibit higher levels of tissue remodeling genes. Most interestingly, these dedifferentiated early postnatal myofibroblasts re-express SMA upon exposure of the adult lung to hypoxia or the pro-fibrotic drug bleomycin. However, unlike during alveolarization, these cells that re-express SMA do not proliferate with hypoxia. In sum, dedifferentiated early postnatal myofibroblasts are a previously undescribed cell type in the adult lung and redifferentiate in response to injury.

## Introduction

During lung development, ∼90% of the gas exchange surface area forms through alveolarization^1^. In altricial species, such as humans and mice, alveolarization is predominantly a postnatal phenomenon and is bi-phasic, consisting of bulk (i.e., classical) and continued alveolarization^1–4^. In mice, bulk alveolarization is rapid, beginning at postnatal day (P) 3 and continuing until P14. This rapid alveolarization occurs through septal eruption of the pre-existing immature septae formed during branching morphogenesis^4,5^. Conversely, continued alveolarization progresses slowly from P14 - P36 by the lifting off of new septae from mature pre-existing septae^1,4,6,7^. During bulk alveolarization, elongated alpha-smooth muscle actin (SMA)^+^ cells, called myofibroblasts, secrete elastin and collagen fibers in response to cues originating primarily from the adjacent epithelium, thereby generating the secondary septae to form alveoli^5,6,8–10^. SMA, encoded by the *Acta2* gene, is the most widely accepted marker for myofibroblasts and additional genes, such as *Aspn, Mustn1, Hhip and Tgfbi,* have recently been described as markers of this population as well based on transcriptomic data^11,12^; however, for these latter genes, validation in terms of protein expression is lacking.

Platelet-derived growth factor receptor (PDGFR)-α^+^ mesenchymal cells are widely implicated as progenitors of myofibroblasts that express smooth muscle cell (SMC) markers SMA and SM22α, and signaling through the PDGFA-PDGFR-α axis is essential for alveolarization^8,9,13–16^. While PDGFR-α^+^ progenitor cell differentiation is critical for rapid generation of myofibroblasts during bulk alveolarization, it is not established whether proliferation of myofibroblasts plays a role in this process. In the context of bleomycin-induced lung fibrosis, our recent findings indicate that differentiated myofibroblasts are proliferative^17^. Moreover, in PDGFR-α^+^ cells isolated from the normal lung at P4, there is a positive correlation between expression levels of SMA and the proliferation marker Ki67^(ref. 18)^. Importantly, after bulk alveolarization at ∼P15, SMA^+^ lung myofibroblasts are close to undetectable^9^. Whether alveolar myofibroblasts undergo apoptosis prior to P15 or dedifferentiate and persist to adulthood remains controversial^15,19–22^. PDGFR-α marks SMA^-^ myofibroblast progenitors in the embryonic and early postnatal lung and SMA^+^ myofibroblasts during bulk alveolarization^9,15,16,18^. Li and colleagues report that following lineage-labeling PDGFR-α^+^ cells between P1-P20, marked cells in the P40 lung are SM22α^-^ and thereby, suggest that the PDGFR-α^+^ lineage persists in the adult lung in a dedifferentiated state^15^. On the other hand, a study utilizing *Fgf18-CreER^T^*^2^ to label early postnatal myofibroblasts indicates apoptosis and clearance of these mesenchymal cells after alveolarization^19^. In support of this finding, a previous study shows substantial apoptosis of fibroblasts isolated from the rat lung after bulk alveolarization^22^.

Herein, our murine studies demonstrate that SMA^+^ myofibroblasts are proliferative during bulk alveolarization, and that at the end of the second postnatal week, the majority of myofibroblasts downregulate SMC markers and then undergo apoptosis. However, a small percentage of the cells that downregulate SMC markers persist to adulthood. Single cell RNA sequencing (scRNA-seq) of these persistent dedifferentiated cells reveals a transcriptomic signature that is unique from previously described lung fibroblasts. Interestingly, the dedifferentiated postnatal myofibroblasts in adult lungs, re-express SMA following hypoxia exposure or bleomycin-induced injury. Thus, we suggest that these dedifferentiated cells are a reserve population in the adult lung that is responsive to injury and may play an important role in subsequent lung remodeling.

## Results

### Early postnatal myofibroblasts accumulate and proliferate during bulk alveolarization

Alveolarization begins with septal eruption at P4 in the mouse lung^4^. Myofibroblasts, elongated SMA^+^ cells in the lung parenchyma distinct from vascular and airway SMCs, are the primary cell type implicated in laying down elastin in the alveolar septa during postnatal alveolarization^8,9,23,24^. To study the dynamics of SMA^+^ myofibroblast accumulation during alveolar septation, lungs were harvested from wild type mice at discrete time points in the early postnatal period until P30.5. Cryosections were stained for SMA and nuclei (DAPI). At P0.5, SMA expression was restricted to SMCs, and myofibroblasts were essentially not detectable in the lung parenchyma (Fig. 1a, b). SMA^+^ myofibroblasts initially appear at P2.5, markedly increase in number until they peak at P11.5, then subsequently decrease rapidly with less than half of the peak number of SMA^+^ myofibroblasts observed at P14.5 and only very rare myofibroblasts discernible by P18.5 (Figs. 1a, b, S1).

**Figure 1.**
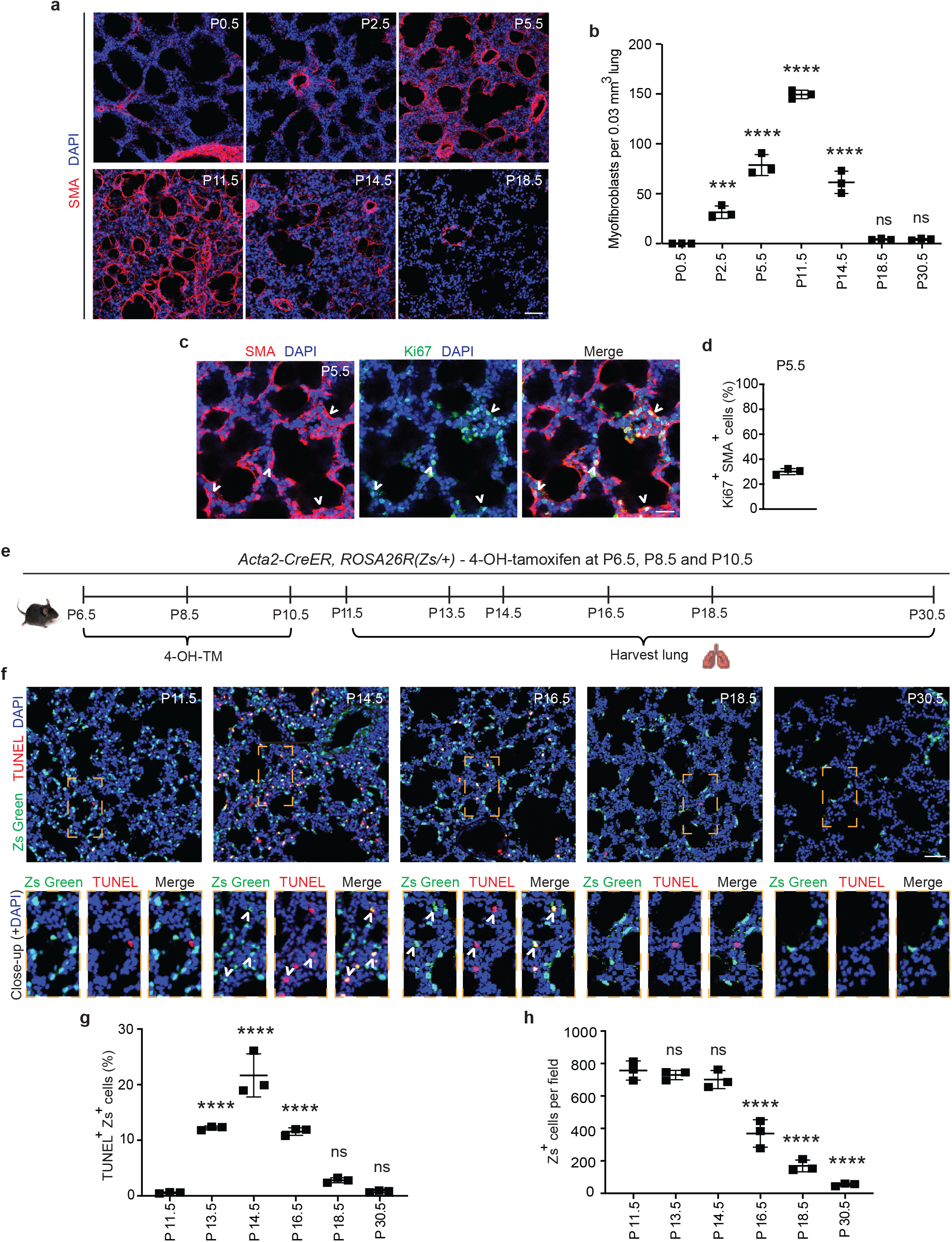
Myofibroblasts proliferate and then undergo apoptosis during bulk alveolarization. Lungs were harvested from postnatal mice at indicated ages, cryosectioned, and stained. **a, c,** Sections from wild type mice were stained for SMA and in **c,** for Ki67 as well. **b**, The number of SMA^+^ myofibroblasts per 0.03 mm^3^ lung volume was quantified from sections as in **a**; n=3 mice per time point. **d**, Percent of SMA^+^ myofibroblasts that were Ki67^+^ was quantified, n=3 mice, 5 sections per mouse and an average of 165 myofibroblasts per section. **e-h**, *Acta2-CreER^T^*^2^, *ROSA26R*(Zs/+) mice were induced with 4-hydroxy-tamoxifen (4-OH TM) at P6.5, P8.5 and P10.5. Schematic is displayed in **e.** In **f,** lung cryosections were stained with the TUNEL assay and imaged for Zs (fate marker). Close-ups of boxed regions are shown below. In **g, h,** the percent of Zs^+^ cells that are TUNEL^+^ and the number of Zs^+^ cells per microscopic field were quantified, respectively. n=3 mice per time point, 3 sections per mouse and an average of 14-271 Zs^+^ cells per section were analyzed. One-way ANOVA with Tukey’s multiple comparison test was used with statistical significance vs. P0.5 in **b** and vs. P11.5 in **g, h**. ns,***, **** indicate not significant, p<0.001, p<0.0001, respectively. Scale bars, 50 μm **(a, f)**, 25 μm **(c)**.

PDGFR-α^+^ progenitor cells have been shown to differentiate into postnatal SMA^+^ myofibroblasts in the early postnatal lung^14–16^. Although, proliferation and differentiation are often inversely correlated^25^, prior studies suggest that differentiated (i.e., SMA^+^) myofibroblasts are proliferative^17,26^. To determine whether SMA^+^ lung myofibroblasts proliferate during bulk alveolarization, cryosections of lungs from wild type mice at P5.5 were stained for SMA and the proliferation marker Ki67 (Fig. 1c). At this time point, 30±2% of SMA^+^ myofibroblasts are Ki67^+^ (Fig. 1d), suggesting that myofibroblast accumulation during bulk alveolarization results from myofibroblast proliferation, in addition to progenitor cell differentiation as indicated by earlier studies^15,16^.

### Majority of early postnatal myofibroblasts dedifferentiate and undergo apoptosis

The reason for the disappearance of almost all SMA^+^ myofibroblasts following bulk alveolarization is controversial. While some studies suggest apoptosis^19–22^, others propose their persistence after downregulation of SMC markers^15^. To evaluate this issue in wild type mice, we immunostained lung sections at P11.5 and P14.5 for the apoptotic marker caspase-3.

Surprisingly, although the lung has numerous caspase-3^+^ apoptotic cells at P14.5, caspase- 3^+^SMA^+^ myofibroblasts were rarely detectable (Fig. S2a). Thus, we postulated that myofibroblasts downregulate SMA prior to undergoing apoptosis. To test this possibility, *Acta2- CreER^T^*^2^*, ROSA26R*(Zs/+) mice were induced with 4-hydroxy-tamoxifen (4-OH-TM) on P6.5, P8.5 and P10.5 to permanently label SMA^+^ myofibroblasts and their lineage (Fig. 1e). Note that, as *Acta2-CreER^T^*^2^ utilizes an inducible Cre-loxP recombination system, expression of the lineage marker (in this manuscript, ZsGreen1 [Zs] or GFP) is retained permanently in lineage^+^ cells even after expression of *Acta2* (encoding SMA) is downregulated. Lung cryosections at specific timepoints from P11.5 - P30.5 were stained with terminal transferase dUTP-digoxygenin nick end-labeling (TUNEL) and DAPI and directly imaged for Zs (Fig. 1f). At P11.5, very rare Zs^+^ cells are TUNEL^+^ (0.6±0.1%; Fig. 1g). Thereafter, there is a significant increase in this apoptotic cell percentage, peaking at P14.5 (21.6±3.9%) and subsequently decreasing such that only rare Zs^+^ cells are observed at P30.5 (Fig. 1h). Although there is a significant reduction in the SMA^+^ myofibroblasts between P11.5 - P14.5, during this time period, the number of Zs^+^ cells are unchanged, and the majority of TUNEL^+^Zs^+^ cells are SMA^-^ (Figs. 1g, S2b, c) validating our hypothesis that dedifferentiation precedes apoptosis. Additionally, we noted Zs^+^ signal within CD68^+^ macrophages, suggesting the clearance of remnants of post-apoptotic myofibroblasts by macrophages (Fig. S3)^19^. Taken together, these findings suggest that at the end of bulk alveolarization, the majority of early postnatal myofibroblasts downregulate SMA and then undergo apoptosis with the accumulated cell debris being cleared by macrophages.

### Select early postnatal myofibroblasts dedifferentiate and persist to adulthood

The staining for SMA and TUNEL and the fate mapping of SMA^+^ cells (see Fig. 1) suggest the presence of rare cells derived from the SMA^+^ population that are TUNEL^-^ and likely SMA^-^ in the lung parenchyma at P30.5. Hence, we next sought to determine whether some of the early postnatal SMA^+^ myofibroblasts dedifferentiate, and their lineage persists into adulthood. To this end, *Acta2-CreER^T^*^2^*, ROSA26R*(mTmG/+) pups were injected with 4-OH-TM at P6.5, P8.5 and P10.5, and lungs were harvested either at P11.5 to assess labeling efficiency, or at P60.5 for lineage tracing (Fig. 2a). Analysis of labeling efficiency indicates that 82±1% of SMA^+^ myofibroblasts are GFP^+^ at P11.5 (Fig. S4). SMA^+^ myofibroblasts largely undergo apoptosis following bulk alveolarization, and at P60.5, rare GFP^+^ cells persist in the lung (Fig. 2b, c). Indeed, ∼9% of the total number of GFP^+^ lung cells at P11.5 are present in the adult lung. At P60.5, of these persisted GFP^+^ cells, 85±4% are SMA^-^ and the rest are SMA^+^ (Fig. 2b, c). These findings suggest that, after bulk alveolarization, >90% of the myofibroblasts undergo apoptosis but of the remaining myofibroblasts which do persist into adulthood, the vast majority are dedifferentiated.

**Figure 2.**
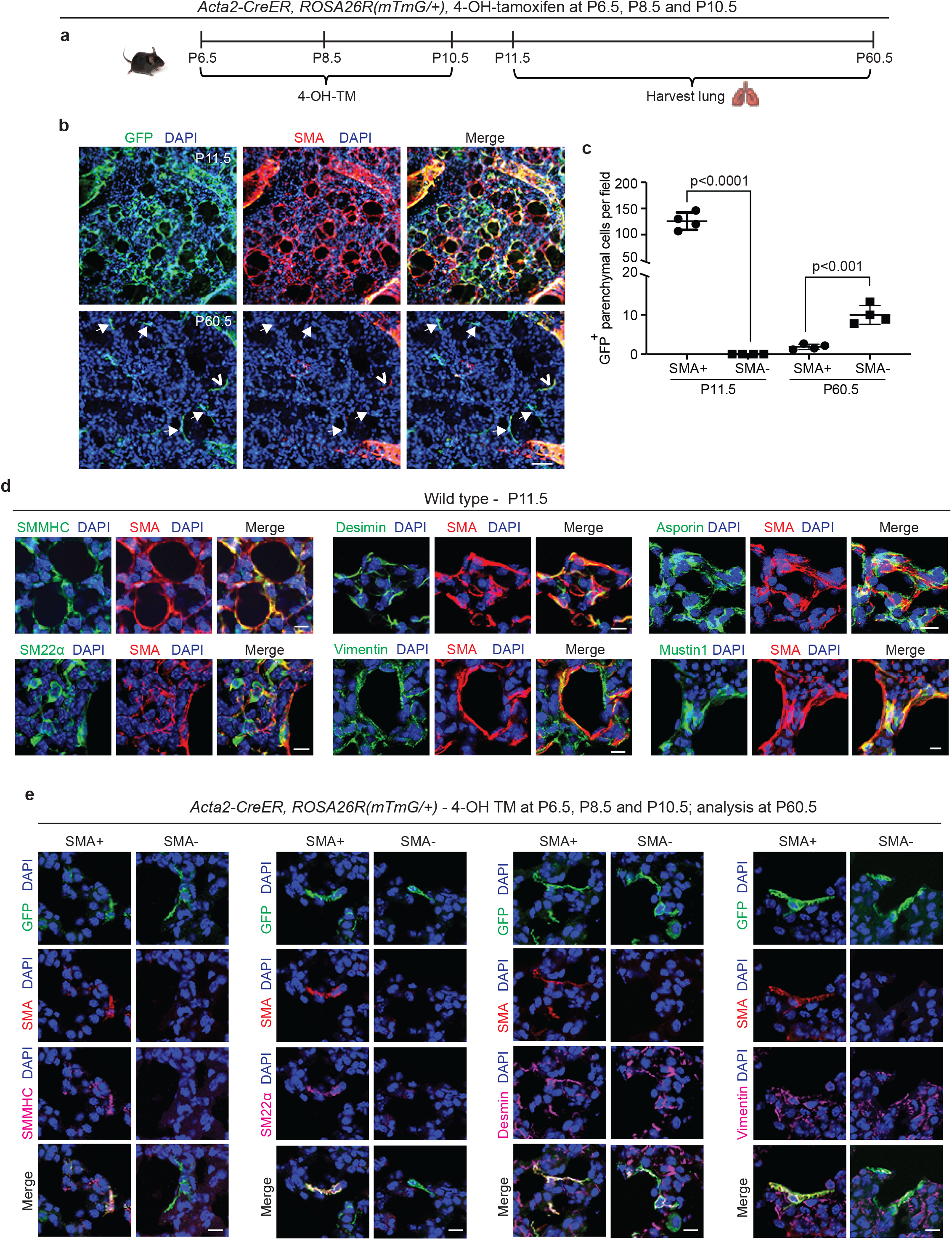
Select SMA^+^ myofibroblasts downregulate smooth muscle cell markers and survive to adulthood. **a-c,** *Acta2*-*CreER^T^*^2^, *ROSA26R*(mTmG/+) mice were induced with 4-OH TM at P6.5, P8.5 and P10.5, and lungs were harvested either at P11.5 or P60.5. Schematic is shown in **a.** In **b,** lung vibratome sections were stained for SMA, GFP (fate marker) and nuclei (DAPI). Open arrowhead indicates GFP^+^SMA^+^ myofibroblast, and arrows with closed heads indicate GFP^+^SMA^-^ dedifferentiated myofibroblasts. In **c**, the number of GFP^+^ cells per microscopic field that are SMA^+^ or SMA^-^ are quantified at P11.5 and P60.5; n=4 mice per time point, 6 sections per mouse and on average 126 (for P11.5) or 5 (for P60.5) GFP^+^ cells were quantified per section. Two-tailed Student’s t-test was used. **d,** Lungs were harvested from wild type mice at P11.5, and parenchymal cryosections were stained for SMA, nuclei (DAPI) and other markers as indicated; n=6. **e,** *Acta2*-*CreER^T^*^2^, *ROSA26R*(mTmG/+) mice were induced with 4-OH TM at P6.5, P8.5 and P10.5, and lungs were harvested at P60.5. Lung vibratome sections were stained for SMA, GFP and nuclei (DAPI) and other markers as indicated. Examples of both SMA^+^ and SMA^-^ cells are shown; n=4-6. Scale bars 50 μm **(b)**, 10 μm **(d, e)**.

Next, we evaluated the gene expression pattern of early postnatal SMA^+^ myofibroblasts at the time of their peak abundance. For this analysis, *Acta2-CreER^T^*^2^*, ROSA26R*(mTmG/+) mice, which received 4-OH-TM at P6.5, P8.5 and P10.5, or wild type mice, were euthanized at P11.5. Lung cryosections were stained for SMA, nuclei (DAPI), cell population-specific markers, and, in the case of *Acta2-CreER^T^*^2^*, ROSA26R*(mTmG/+) mice, for GFP as well (Figs. 2d, S5a) . At P11.5, SMA^+^ myofibroblasts express other SMC markers, including, smooth muscle myosin heavy chain (SMMHC) and transgelin (SM22α), and mesenchymal markers, desmin and vimentin (Fig. 2d). Known markers of SMA^+^ myofibroblasts, such as asporin, mustin1 and transforming growth factor beta-induced (TGFBI)^11,12,27^, are also expressed (Figs. 2d, S5a). However, markers of lipofibroblasts (adipose differentiation related protein [ADRP]), macrophages (CD68, sialic acid-binding immunoglobulin-like lectin F [Siglec-F]) and epithelium (prosurfactant protein C [proSPC]) were not detected (Fig. S5a).

Additionally, the expression pattern of the lineage derived from early postnatal myofibroblasts that persist to adulthood was analyzed. *Acta2-CreER^T^*^2^*, ROSA26R*(mTmG/+) mice were injected with 4-OH-TM on P6.5, P8.5 and P10.5, and immunohistochemistry of lung sections at P60.5 was performed to assess marker expression in GFP^+^ cells that were also SMA^+^ or SMA^-^ (Fig. 2e). SMMHC and SM22α track with SMA expression. In contrast, both GFP^+^SMA^+^ and GFP^+^SMA^-^ cell types express desmin, vimentin, mustin1 and TGFBI but do not express CD68 nor ADRP (Figs. 2e, S5b). According to our previous study, PDGFR-β is expressed in different subtypes of healthy lung fibroblasts^17^, and thus, we queried whether GFP^+^SMA^+^ and GFP^+^SMA^-^ cell populations express PDGFR-β. Both cell populations exhibit heterogeneity in terms of PDGFR-β expression, such that only some of the SMA^+^GFP^+^ cells and SMA^-^GFP^+^ cells are PDGFR-β^+^ (Fig. S6).

### Single cell transcriptomic analysis of the lineage-traced, early postnatal myofibroblasts in adulthood

To evaluate the transcriptome of the lineage of early postnatal myofibroblasts in the adult lung, lineage labeled cells were analyzed by scRNA-seq. *Acta2-CreER^T^*^2^*, ROSA26R*(Zs/+) mice were injected with 4-OH TM on P6.5, P8.5 and P10.5 to label SMA^+^ cells with Zs (Fig. 3a). Mice were aged to P60.5 and euthanized, and lungs were harvested. Single cell lung suspensions were stained with DAPI, and Zs^+^DAPI^-^ live cells were isolated with fluorescence activated cell sorting (FACS). Isolated cells were then subjected to droplet-based single-cell sequencing (DropSeq), and transcriptomic data was processed with Cell Ranger v3. Mouse transcriptome mm10 modified with the addition of the Zs gene sequence was used as the reference genome.

**Figure 3.**
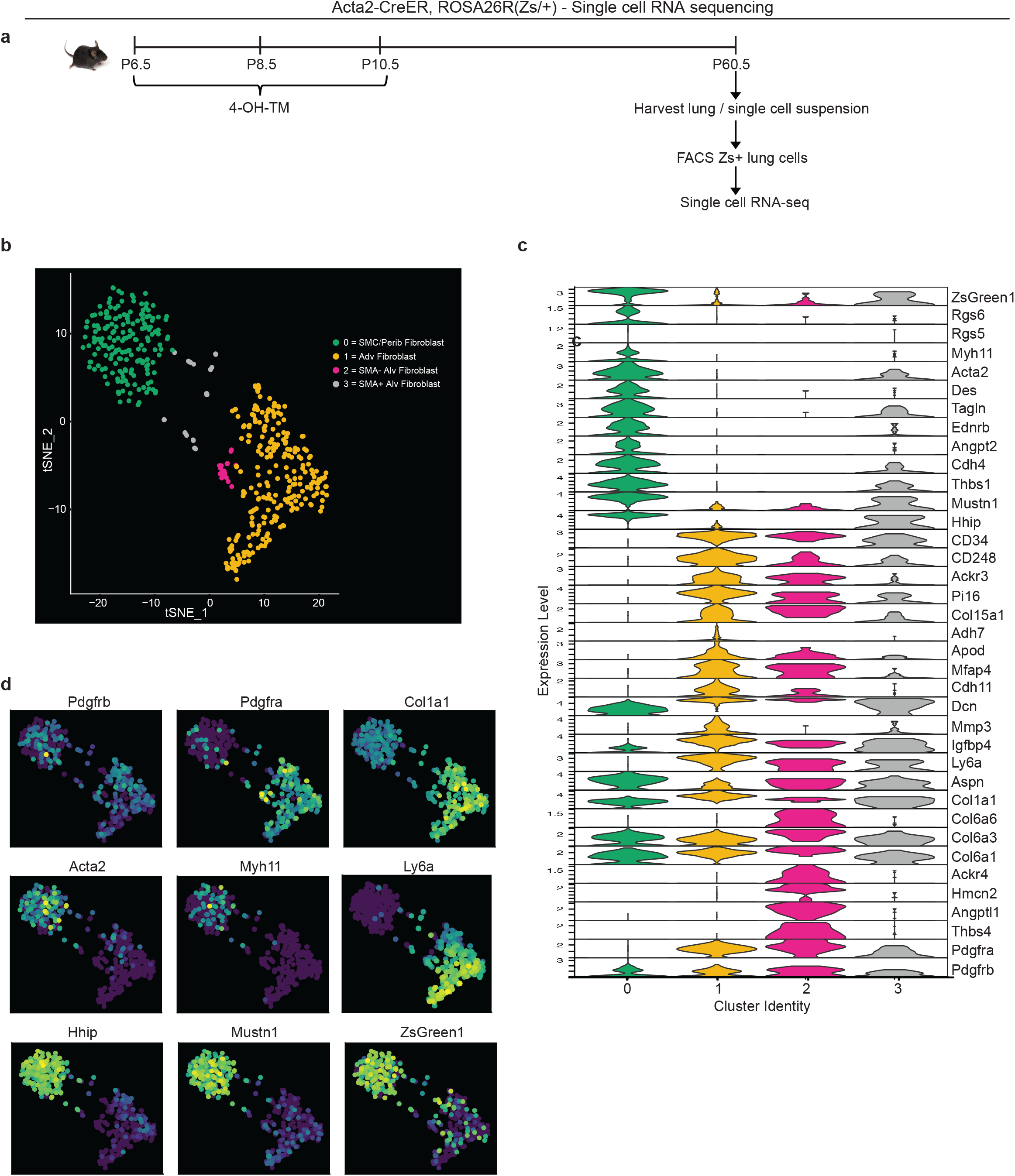
Single cell RNA sequencing of lineage-traced early postnatal SMA^+^ cells in adult lung. *Acta2-CreER^T^*^2^*, ROSA26R*(Zs/+) mice were induced with 4-OH TM at P6.5, P8.5 and P10.5. At P60.5, lungs were harvested. Zs^+^ cells of the lung were isolated by FACS and subjected to scRNA library construction (10X Genomics), sequencing, annotation and clustering. **a**, Schematic of experiment. **b**, tSNE plot with cells colored by their categorized cell-type identity. **c**, Violin plot showing expression levels of representative marker genes in each cluster. **d**, Same tSNE plot as in **b**, colored based on normalized expression levels of indicated transcripts.

Cell annotation and clustering were performed based on the cell-specific transcriptional marker gene expression as described by Tsukui et al.^28^ T-distributed stochastic neighbor embedding (t-SNE) was performed to visualize gene expression, and four clusters were defined (Fig. 3b). Cluster 0 is composed of SMCs and peribronchial fibroblasts with the highest expression of SMC markers (*Acta2, Tagln, Myh11* and *Des*) and peribronchial fibroblast markers (*Hhip* and *Aspn*; Fig. 3c, d). As expected, this cluster exhibits low expression of *Pdgfra.* Adventitial fibroblasts are represented by cluster 1 with high *Pdgfra, Ly6a* and *Col1a1* (Fig. 3c, d). This cluster is also enriched in *Col15a1* and *Adh7*.

Clusters 2 and 3 are alveolar fibroblasts with high expression of *Pdgfra*, *Aspn* and *CD34* (Fig. 3c). We identify cluster 3 as the rare cells of the lineage of early postnatal alveolar myofibroblasts that have remained SMA^+^ and with representative cluster identity of *Acta2^+^Tagln^+^Pdgfra^+^Col6a3^+^Hhip^+^Mustn1^+^Dcn^+^* and cluster 2 as the lineage of early postnatal alveolar myofibroblasts that have downregulated SMA and with gene expression signatures of *Pdgfra^+^Col6a3^+^Col6a6^+^Acta2^-^Tagln^-^Hhip^-^.* Interestingly, cluster 2, but not cluster 3, is highly enriched in *Thbs4, Angptl1, Hmcn2 and Ackr4* (Fig. 3c), genes known to be involved in tissue remodeling^29–32^. We therefore next examined whether these SMA^-^ dedifferentiated cells derived from early postnatal myofibroblasts contribute to adult pathologies with substantial lung remodeling.

### Dedifferentiated early postnatal SMA^+^ cells contribute to hypoxia-induced myofibroblast accumulation

Although SMA^+^ alveolar myofibroblasts are quite rare in the normal adult lung, they accumulate with exposure of mice to hypoxia^33,34^. Thus, we interrogated whether dedifferentiated early postnatal SMA^+^ cells re-express SMA and contribute to the pathological lung myofibroblast pool in hypoxic adult mice. To this end, *Acta2-CreER^T^*^2^*, ROSA26R*(mTmG/+) mice were induced with 4-OH-TM on P6.5, P8.5 and P10.5, and then rested until P60.5, after which they were subjected to 21 days of normoxia or hypoxia to induce pulmonary hypertension and right ventricle hypertrophy (Fig. 4a-c). Twelve hours prior to euthanasia, mice were injected - intraperitoneally with 5-ethynyl-2’-deoxyuridine (EdU) to assess cellular proliferation. In the adult lung after hypoxia exposure, 31±4% of the SMA^+^ myofibroblasts are GFP^+^ (Fig. 4d, e), suggesting that ∼one-third of hypoxia-induced SMA^+^ myofibroblasts originate from redifferentiation of early postnatal myofibroblasts. Additionally, of the total GFP^+^ cells in the parenchyma of the adult lung (excluding SMCs), 96±3% are SMA^+^ after hypoxia for 21 days as compared to 14±2% under normoxic conditions (Fig. 4f). Interestingly, GFP^+^ alveolar cells were not proliferative under hypoxic conditions (Fig. 4g, h). Thus, a significant number of lineage^+^ SMA^-^ cells contribute to hypoxia-induced SMA^+^ lung myofibroblasts predominantly by differentiation with limited, if any, proliferation.

**Figure 4.**
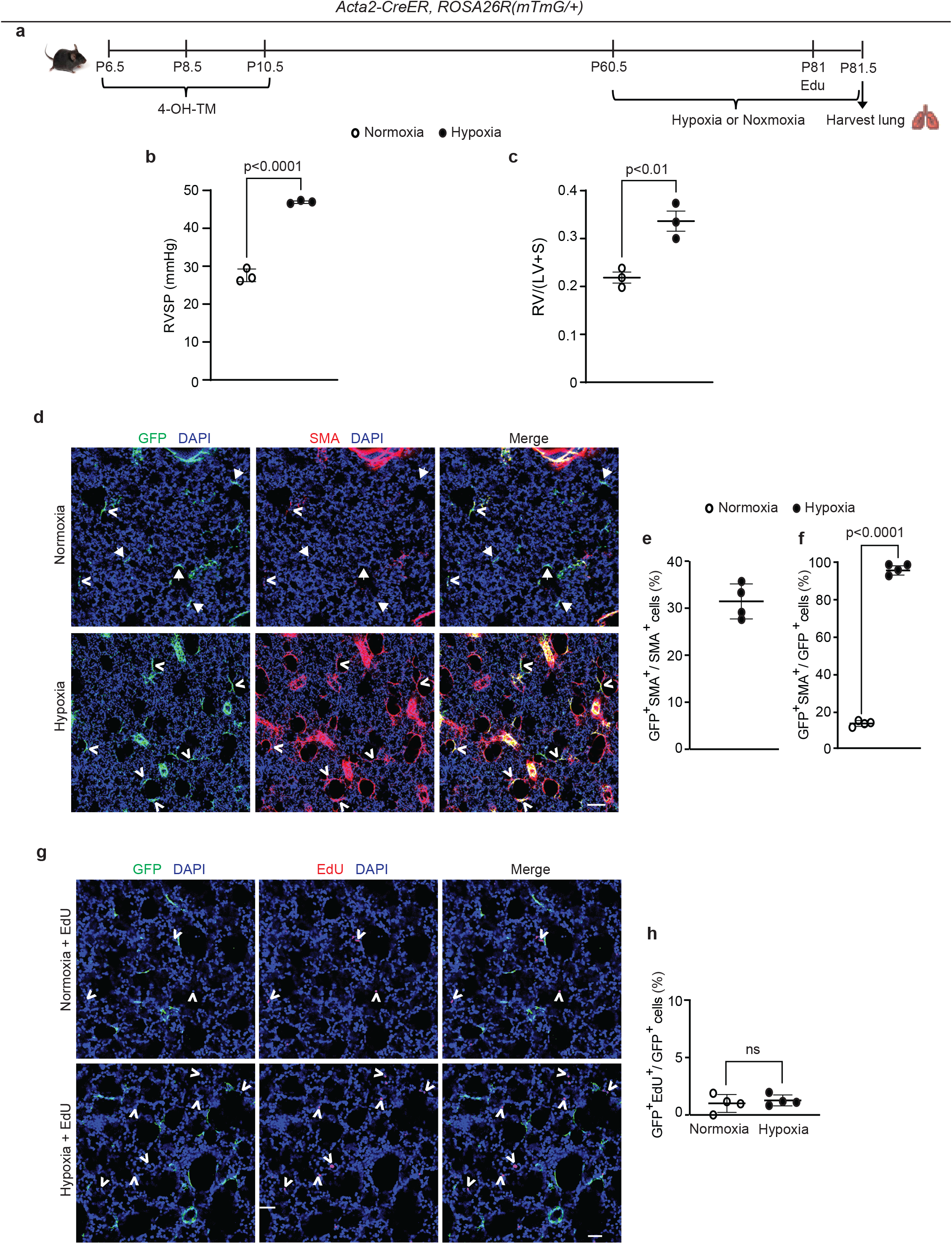
Dedifferentiated early postnatal SMA^+^ cells redifferentiate upon hypoxia exposure in the adult lung. *Acta2*-*CreER^T^*^2^, *ROSA26R*(mTmG/+) mice were induced with 4-OH TM at P6.5, P8.5 and P10.5, rested until P60.5, exposed to hypoxia (FiO_2_ 10%) or normoxia for 21 days, subjected to right ventricular systolic pressure (RVSP) measurements and then euthanized. Twelve hours prior to euthanasia, mice were injected with EdU. **a**, Schematic of experimental set up. **b,** RVSP was quantified as indicated. **c,** The ratio of the weight of the right ventricle (RV) to that of the sum of left ventricle (LV) and septum (S) was measured. n=3 mice for each group. **d**, Lung cryosections were stained for SMA, GFP (fate marker) and nuclei (DAPI). n=4 mice per group. Open arrowheads indicate GFP^+^SMA^+^ cells and arrows with closed heads indicate GFP^+^SMA^-^ cells. **e**, Percentage of SMA^+^ cells that are GFP^+^ was quantified; n=4 mice, 5-8 sections per mouse, an average of 57 SMA^+^ cells were quantified per section. **f**, Percentage of GFP^+^ cells that are SMA^+^ was quantified; n=4 mice, 5-8 sections per mouse, an average of 13 (normoxia) or 18 (hypoxia) GFP^+^ cells were quantified per section. Two-tailed Student’s t-test was performed. **g,** Lung cryosections were stained for EdU, GFP (fate marker) and nuclei (DAPI). Open arrowheads indicate EdU^+^ cells. **h**, The percent of GFP^+^ cells that are EdU^+^ is quantified. For each treatment group, n=4 mice, 6 sections per mouse, average 15 (normoxia) or 16 (hypoxia) GFP^+^ cells per section quantified; ‘ns’ indicates not significant. Scale bars, 100 μm **(d)**, 50 μm **(g)**.

### Dedifferentiated early postnatal SMA^+^ cells redifferentiate during bleomycin-induced lung fibrosis

To address whether this phenomenon is specific to hypoxia, the bleomycin lung injury model was next used to study whether lineage^+^ cells contribute to accumulation of SMA^+^ myofibroblasts and fibrosis in the lung^35^. Most of these SMA^+^ myofibroblasts form discrete thick interstitial patches in the lung (herein, referred to as interstitial myofibroblasts and the regions they occupy as fibrotic); however, some of the SMA^+^ myofibroblasts are located in relatively normal appearing non-fibrotic alveolar regions (herein, these myofibroblasts are referred to as alveolar myofibroblasts and the regions they occupy as intact) and have an elongated morphology that is similar to alveolar SMA^+^ myofibroblasts in the normal lung (Fig. 5).

**Figure 5.**
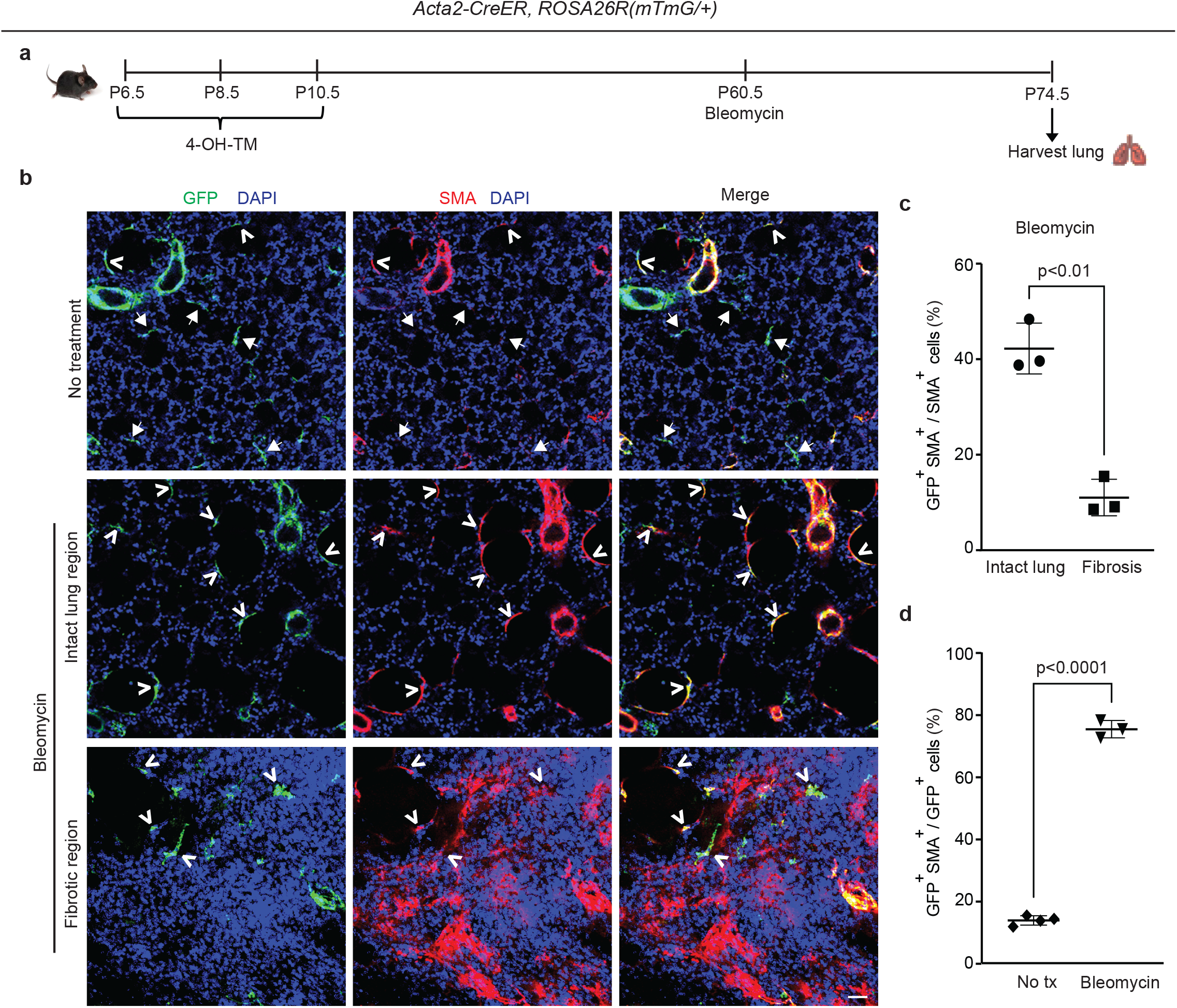
Dedifferentiated early postnatal SMA^+^ cells redifferentiate with bleomycin- induced injury in the adult lung. *Acta2*-*CreER^T^*^2^, *ROSA26R*(mTmG/+) were induced with 4-OH TM at P6.5, P8.5 and P10.5 and then treated or not treated with a single orotracheal dose of bleomycin at P60.5. After 14 days, mice were euthanized, and lungs were harvested. **a**, Schematic of experimental set up. **b**, Lung cryosections were stained for SMA, GFP (fate marker) and nuclei (DAPI). Open arrowheads indicate GFP^+^SMA^+^ cells, and arrows with closed heads indicate GFP^+^SMA^-^ cells. Scale bar, 50 μm. **c**, Percentage of SMA^+^ cells that are GFP^+^ was quantified in ‘intact lung region’ and ‘fibrotic region’ in the bleomycin-treated group; n=3 mice, 5 sections per mouse, an average of 33 (intact lung region) or 175 (fibrotic region) SMA^+^ cells were quantified per section. **d**, Percentage of GFP^+^ cells that are SMA^+^ was quantified in not treated and bleomycin group; n=3-4 mice, 5 sections per mouse, an average of 13 (bleomycin) or 14 (not treated) GFP^+^ cells were quantified per section. Two-tailed Student’s t- test was used in **c** and **d**.

Previous studies from our group and others reported that pre-existing PDGFR-β^+^ cells are the primary source of bleomycin-induced SMA^+^ interstitial myofibroblasts, whereas adult SMA^+^ cells provide a limited (maximum of ∼10%) contribution^17,36^. Additionally, some of the dedifferentiated elongated alveolar cells in the adult that derive from early SMA^+^ cells are PDGFR-β^+^ (see Fig. S6). To investigate whether cells in the adult lung parenchyma that derive from early postnatal SMA^+^ myofibroblasts give rise to pathological myofibroblasts during fibrosis, *Acta2-CreER^T^*^2^*, ROSA26R*(mTmG/+) mice were induced with 4-OH-TM on P6.5, P8.5 and P10.5 and then were or were not subjected to a single dose of orotracheal bleomycin at P60.5 (Fig. 5a). Fourteen days later, lungs were harvested, and sections were stained for SMA, GFP and nuclei (DAPI) and quantified (Fig. 5a-c). After bleomycin exposure, in non-fibrotic intact lung regions, 42±5% SMA^+^ myofibroblasts were GFP^+^ cells, whereas in fibrotic regions, among interstitial myofibroblasts this contribution was 11±4%, which is similar to that seen following adult *Acta2* lineage labeling^17^ (Fig. 5b, c). Furthermore, while only 14±1% of GFP^+^ cells are SMA^+^ in the untreated normal lung, 75±3% of GFP^+^ cells are SMA^+^ in the intact lung regions of the bleomycin-treated fibrotic lung (Fig. 5d). Taken together, these findings suggest that the majority of persistent early postnatal dedifferentiated myofibroblasts redifferentiate during bleomycin-induced lung fibrosis to contribute to SMA^+^ alveolar myofibroblasts in the intact lung regions, while not contributing to patchy fibrotic regions.

## Discussion

In this study, we describe the dynamics of alveolar myofibroblast accumulation and gene expression in the normal mouse lung during and after bulk alveologenesis and how, in adulthood, this lineage responds to injury (Fig. 6). SMA^+^ myofibroblasts initially appear in the lung at ∼P2.5, proliferate and rapidly accumulate, with their numbers peaking at ∼P11.5 (Fig. 1). Consistent with previous reports^6,15,19,22^, over the next several days, these numbers are rapidly reduced. A group of studies suggest that myofibroblasts undergo apoptosis at the tail end of bulk alveolarization^19–22^. For instance, tamoxifen-induction of *Fgf18-CreER^T^*^2^*, ROSA26R-tdTomato* mice at P5-P8 demonstrates that tdTomato^+^ alveolar myofibroblasts are cleared after the initial phase of rapid alveologenesis^19^. In contrast, the authors of a distinct study suggest the persistence of PDGFR-α^+^ myofibroblasts into adulthood after downregulation of SMC markers^20^. In this latter study, the lineage of PDGFR-α^+^ cells were marked by daily injection of doxycycline from P1-P20, and lungs were analyzed at P40. The presence of lineage marked cells that are almost exclusively SM22α^-^ at P40 was interpreted to indicate that PDGFR-α^+^SM22α^+^ cells downregulate SM22α and persist^15^. An alternative explanation is the following: i) PDGFR- α^+^SM22α^+^ myofibroblasts undergo apoptosis by ∼P15; and ii) PDGFR-α^+^SMC marker^-^ cells are labeled during the P15-20 time period, and cells of this lineage are present at P40. In light of this issue, we induced *Acta2-CreER^T^*^2^, *ROSA26R*(mTmG/+) mice with tamoxifen at P6.5, P8.5 and P10.5 to label and fate map cells that express SMA, the most widely accepted myofibroblast marker (Fig. 2). Our results indicate that the vast majority of lineage marked cells downregulate SMA expression and then undergo apoptosis but a small minority of GFP^+^ cells persist. Indeed, ∼9% of the total number of lineage marked lung cells at P11.5 are present in the adult lung and 90% of these cells are dedifferentiated (i.e., SMA^-^).

**Figure 6.**
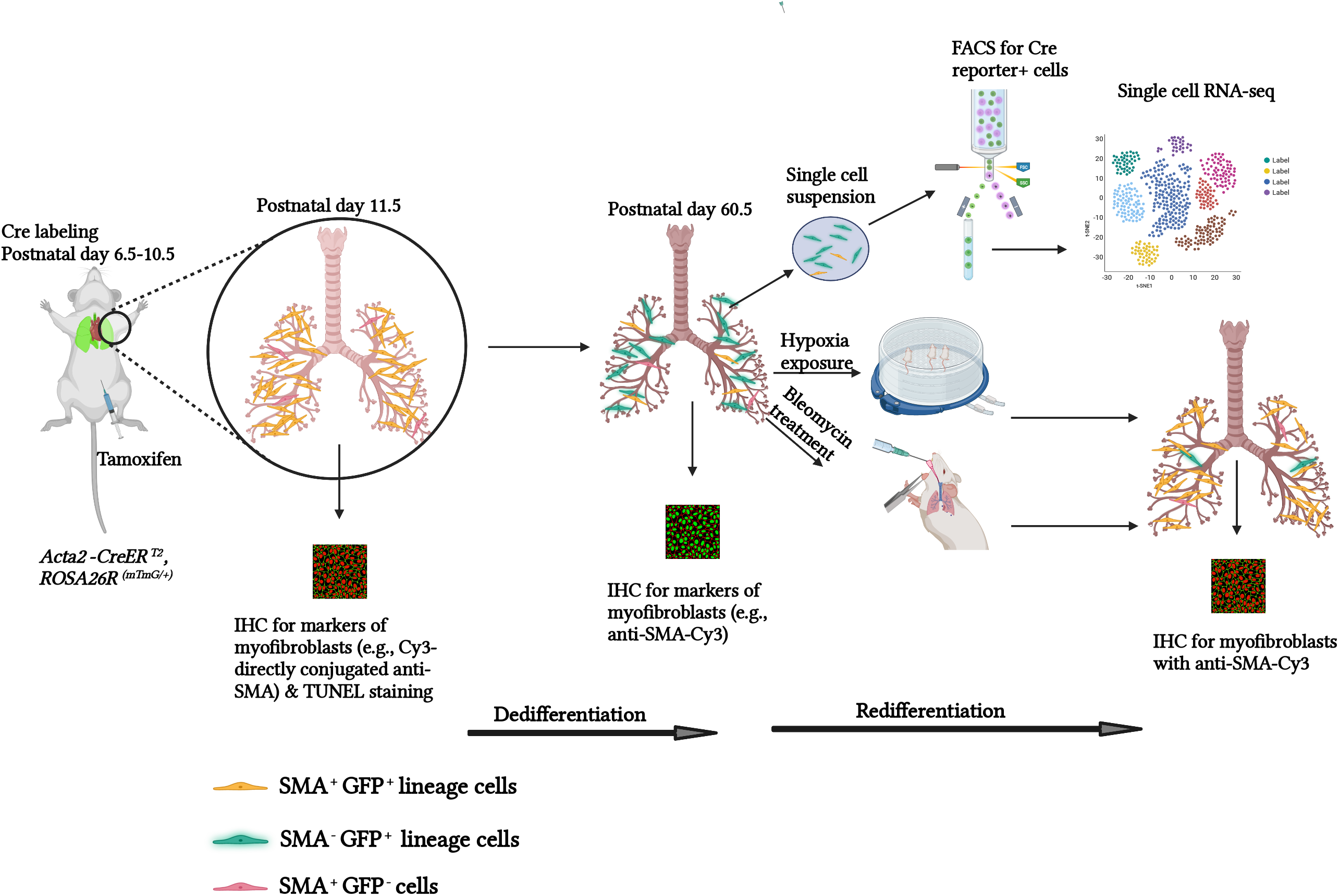
Schematic representation of experiments and findings. Tamoxifen was administered to *Acta2-CreER, ROSA26R*(mTmG/+) pups at P6.5, P8.5 and 10.5 to label lung myofibroblasts by P11.5. Most of these myofibroblasts downregulate SMA by the end of bulk alveolarization (∼P14.5) and then undergo apoptosis whereas a minority persist into adulthood. These SMA^-^GFP^+^ cells are an important source of SMA^+^ lung myofibroblasts in adults exposed to hypoxia or bleomycin disease, suggesting that they may play a pathological role in adult lung disease. Created with BioRender.

Using *Acta2-CreER^T^*^2^, *ROSA26R*(Zs/+) mice and scRNA-seq, the transcriptomes of dedifferentiated (SMA^-^Zs^+^) and differentiated (SMA^+^Zs^+^) adult lung parenchymal cells that derive from the early postnatal SMA^+^ cells were compared (Fig. 3). Cluster 2 comprising dedifferentiated alveolar fibroblasts were characterized by expression of *Pdgfra, CD34* and *Aspn*. *Aspn* encodes asporin which interestingly is implicated in promoting TGF-β-induced lung myofibroblast differentiation^37^. This cluster also expresses *Thbs4, Angptl1, Hmcn2 and Ackr4,* genes reputedly involved in tissue remodeling^29,31,32,38^. For instance, thrombospondin-4 (encoded by *Thsp4*) regulates the production and assembly of collagen, and in the pathological context of cardiac pressure overload, thrombospondin-4 prevents excess ECM deposition and myocardial hypertrophy^39^. In contrast to this dedifferentiated cluster, the SMA^+^ alveolar myofibroblasts of cluster 3 express *Acta2, Tagln, Thbs1* and do not express the group of remodeling genes. Finally, the presence of cluster 1, identified as subclass of adventitial fibroblasts, was unexpected in our dataset of cells derived from early postnatal SMA^+^ cells. We put forth two possibilities for the appearance of this adventitial fibroblast cluster: 1) SMA^+^ adventitial fibroblasts are present in the early postnatal lung and persist; or 2) SMA^+^ cells in the early postnatal lung transdifferentiate into adventitial fibroblasts during postnatal maturation or adulthood.

Our studies indicate that dedifferentiated myofibroblasts redifferentiate to express SMA in the adult lung in response to an altered oxygen environment or bleomycin exposure (Figs. 4, 5). Notably, during hypoxia, the dynamics of SMA expression in these cells is reminiscent of alveolar myofibroblast progenitors during the initial stages of postnatal alveolarization: progenitors start as SMA^-^ and by ∼P2.5, begin to express SMA, coincident with the change in oxygen levels following birth. With bleomycin exposure in the adult, dedifferentiated cells redifferentiate to give rise to myofibroblasts in intact lung regions but not in highly fibrosed areas of the lung. Interestingly, a recent study reports that PDGFR-α^+^ cells are SM22α ^-^ in the adult mouse lung, and bleomycin exposure after labeling this lineage in the adult results in lineage-tagged lung myofibroblasts^15^. Taking this work and our findings together suggests that the PDGFR-α ^+^ adult lung cells which do not derive from early postnatal SMA^+^ alveolar myofibroblasts are likely to contribute to fibrotic foci.

Mechanisms underlying the accumulation of dedifferentiated cell-derived myofibroblasts in intact lung regions during bleomycin exposure are not elucidated, but we propose that the hypoxic environment of fibrotic lung regions may be inductive. Indeed, a hypoxia-inducible factor 1α - pyruvate dehydrogenase kinase 1 signaling axis potentiates transforming growth factor (TGF)-β induced differentiation of fibroblasts to myofibroblasts^40,41^. Additionally, cytokines released during fibrogenesis, such as TGF-β and interleukin-1^(ref. 42–44)^, may transform undifferentiated lung cells to myofibroblasts. Interestingly, under hypoxia, redifferentiated myofibroblasts are essentially not proliferative, which is in contrast to the substantial percentage of Ki67^+^ alveolar myofibroblasts observed during early postnatal development (Figs. 1, 4).

Finally, we speculate that dedifferentiated adult lung cells are specialized to respond to injury. As redifferentiated lineage^+^ cells accumulate in intact lung regions during fibrosis, they likely have a distinct role from that of high collagen-producing myofibroblasts. The significance of this novel cell type that differentiates during adult lung pathologies is yet to be uncovered. The lung undergoes extensive remodeling under hypoxic and fibrotic conditions^45–47^ and given the expression of genes implicated in tissue remodeling in the dedifferentiated cell cluster by scRNA-seq, it will be important in future studies to elucidate the role of redifferentiated myofibroblasts in lung remodeling post-injury.

## Methods

### Animals

All mouse experiments were performed in accordance with ethical regulations of the IACUC at Yale University. C57BL/6 wild type mice were used. *Acta2-CreER^T^*^2^ mice have been described previously^48^. Cre reporters *ROSA26R*(mTmG/mTmG) and *ROSA26R*(ZsGreen1/ZsGreen1) were obtained from Jackson Laboratory^49,50^. ZsGreen1 is abbreviated as Zs. Experiments utilized male and female mice.

### Fate mapping, hypoxia and bleomycin treatment

For fate mapping experiments, *Acta2-CreER^T^*^2^*, ROSA26R*(mTmG/+) mice were injected intraperitoneally on P6.5, P8.5 and P10.5 with 200 μg of 4-OH-TM per day. At P60.5, mice were subjected to either hypoxia or bleomycin treatment. For hypoxia experiments, mice were housed in a rodent chamber with a calibrated oxygen controller and sensor (BioSpherix) and exposed to 10% FiO_2_ (hypoxia) or room air (normoxia control) for 21 days. Right ventricular systolic pressure was measured by inserting a catheter into the right ventricle (RV) via the right jugular vein. Mice were sacrificed, and the lungs and heart were harvested. The weight ratio of the RV/(left ventricle + septum) was assessed as described earlier^33^. Alternatively, to induce lung fibrosis, a single dose of bleomycin (1.5 U/kg body weight) was or was not (control) administered orotracheally. Fourteen days later, mice were euthanized, and lungs were harvested. From *Acta2-CreER^T^*^2^*, ROSA26R*(mTmG/+) mice, lung sections were stained for SMA, GFP and nuclei (DAPI). Contribution of the *Acta2*-*CreER^T^*^2^ lineage to accumulated lung SMA^+^ myofibroblasts in response to hypoxia or bleomycin was determined by scoring the percentage of myofibroblasts (parenchymal elongated SMA^+^DAPI^+^ cells) that expressed lineage marker GFP. *Acta2-CreER^T^*^2^*, ROSA26R*(Zs/+) mice were utilized for scRNA-seq.

### Lung preparation and immunohistochemistry

Mice were sacrificed by isoflurane inhalation, and the pulmonary vasculature was flushed by injecting phosphate buffered saline (PBS) through the RV. For vibratome sectioning, lungs were inflated by infusing 2% low-melting agarose through the trachea with an angiocatheter. Harvested lungs were incubated in ice cold PBS for 30 min followed by Dent’s fixative (4:1 methanol/dimethyl sulfoxide) overnight at 4°C, stored in 100% methanol at -20°C for a minimum of two days. For immunohistochemistry, lungs were bleached in 5% H_2_O_2_ in methanol, followed by rehydration sequentially in 75%, 50% and 25% and 0% methanol in PBS. A vibratome was used to cut 150 μm thick sections. For preparing cryosections, lungs were fixed in 4% paraformaldehyde (PFA) overnight, washed and then incubated in 30% sucrose for at least 3 days. Lungs were then embedded in optical cutting temperature compound (OCT-Tissue Tek), frozen in dry ice and stored at -20°C or -80°C. A cryotome was used to cut 10-30 µm thick sections. For immunohistochemistry, vibratome or frozen sections were blocked in 5% goat serum in PBS containing 0.1% Triton X-100 (PBS-T), washed with PBS-T and incubated with primary antibodies at 4°C overnight. On the next day, sections were washed and incubated in secondary antibodies for 2 h. After washing in PBS-T, sections were mounted in fluorescence mounting medium (DAKO) or glycerol:methanol (1:1) mountant. Glycerol:methanol mountant was used to quench endogenous tomato fluorescence in the cryosections in studies using the mTmG reporter.

Primary antibodies used were chicken anti-GFP (1:100, Abcam), rat anti-CD68 (1:200, Biorad), rabbit anti-desmin (1:200, Abcam), rabbit anti-vimentin (1:200, Abcam), rabbit anti- SM22α (1:500, Abcam), rabbit anti-SMMHC (1:100, Thermo Scientific-Alfa Aesar), rabbit anti- Ki67 (1:100, Invitrogen), rabbit anti-asporin (1:200, Abcam), rabbit anti-mustin1 (1:200, Abcam), rabbit anti-TGFBI (1:200, Abcam), rabbit anti-Pro-SPC (1:200, Abcam), rabbit anti-Siglec-F (1:200, Abcam), rabbit anti-ADRP (1:200, Abcam), directly conjugated Cy3 anti-SMA (1:150 - 1:250, Sigma-Aldrich), and biotinylated anti-PDGFR-β (1:50, R&D). Elite ABC reagents (Vector Laboratories) and fluorescein tyramide system (PerkinElmer) were used to amplify biotinylated PDGFR-β staining as described previously^17^. Secondary antibodies were conjugated to either Alexa-488, −564, −647 fluorophores (1:250-500, Invitrogen). Nuclei were visualized with DAPI (1:1000, Sigma-Aldrich).

### Proliferation of hypoxia-induced myofibroblasts derived from early postnatal SMA^+^ cells

*Acta2-CreER^T^*^2^*, ROSA26R*(mTmG/+) mice were injected with 4-OH-TM (200 µg per day) at P6.5, P8.5 and P10.5, and then starting at P60.5, mice were subjected to hypoxia for 21 days. Twelve hours prior to euthanasia, mice were injected intraperitoneally with 2.5 mg of 5-ethynyl- 2’-deoxyuridine (EdU; Thermo Fisher Scientific). Lungs were harvested, fixed in 4% PFA overnight, permeabilized in 0.5% PBS-T for 30 min and stained with the Click-iT EdU Alexa Fluor Imaging Kit per instructions of the manufacturer (Thermo Fisher Scientific). Sections were co-stained for GFP and nuclei (DAPI). The percent of GFP^+^ cells that express EdU were quantified.

### TUNEL assay

TUNEL assay was performed using ApopTag *in situ* apoptosis detection kit (Millipore- Sigma). Briefly, *Acta2-CreER^T^*^2^*, ROSA26R*(Zs/+) mice were injected with 4-OH TM (200 µg per day) at P6.5, P8.5 and P10.5. Lungs were harvested on P11.5, P13.5, P14.5, P16.5 and P18.5 and P30.5 and fixed in 4% PFA. Cyrosections (10 µm) were treated with pre-cooled ethanol:acetic acid (2:1) for 5 min at -20°C and washed in PBS. Slides were then processed according to instructions of the manufacturer, culminating in incubation with anti-digoxigenin-rhodamine fluorochrome conjugate for 30 min at RT, protected from light. Sections were washed in PBS, mounted and directly imaged for Zs and rhodamine.

### Single-cell RNA sequencing

#### Sample preparation and sequencing

Five *Acta2-CreER^T^*^2^*, ROSA26R*(Zs/+) mice were injected with 4-OH-TM (200 µg per day) at P6.5, P8.5 and P10.5. At P60.5, lungs were collected, minced and incubated at 37°C for 40 min in the enzyme mixture from Miltenyi Biotec Lung Dissociation Kit. During this incubation, tissue was subjected twice to gentle lung dissociation protocol in a gentleMACS dissociator (Miltenyi Biotec). After inhibiting enzymatic activity with 10% FBS, single cell suspensions from each mouse were pooled together and passed through a 100 µm cell strainer and centrifuged at 700 × g for 10 min at 4°C. The pellet was washed in 1X PBS with 1% FBS, DAPI (nuclear stain) added to the cell suspension, and Zs^+^DAPI^-^ cells were isolated by FACS. Trypan blue was used to aid in cell counting. The cells were pooled together in 0.04% bovine serum albumin in PBS and proceeded to library preparation and DropSeq. Construction of single cell 3’ RNA-seq libraries and sequencing were undertaken as described previously^17^. Sequencing data was processed with Cell Ranger v 3.1.0. Mouse transcriptome mm10 that includes the sequence for the gene ZsGreen1 was used as the reference genome.

### Cell barcode clustering and annotation

All analyses were performed in R (version 3.6.1) using the package Seurat (version 3.1.0). UMI counts were scaled to 10,000 UMIs per cell, then natural log transformed with a pseudo-count of one (log((TPM/100) + 1)). Feature selection, principal component analysis, neighbor embedding and Louvain modularity clustering were recursively performed on the data for the purpose of identifying clusters of discretely identifiable cell populations. Clusters were then annotated as either a multiplet population or known cell type by expert curation of transcriptional marker genes and concordance with the literature. Multiplets were identified as cell populations whose transcriptomic profile resembled a combination of two or more cell populations found in the data. Additionally, cells with less than 900 transcripts or greater than 10% mitochondrial transcripts were removed. Non-mesenchymal cell types were then discarded, based on expression hallmarks of epithelial (Epcam, Cdh1), endothelial (Pecam1, Cdh5, Vwf) or immune cells (Ptprc). The remaining 460 cells were used to generate t-SNE plots for visualizing gene expression. For this embedding, the top 1,000 variable genes were selected using Seurat’s FindVariableFeatures implementation under default parameters, these genes were scaled and used for principal component analysis. The top 8 principal components were used with Seurat’s Run TSNE implementation with the seed parameter equal to 7 to generate the figure.

### Imaging

Images were acquired with Leica SP5 or SP8 confocal microscope or PerkinElmer UltraView Vox Spinning Disc confocal microscope. For image processing, analysis and cell counting, Volocity software (PerkinElmer), Adobe Photoshop and Adobe Illustrator were used.

### Statistical analysis

Student’s t-test or ANOVA with Tukey’s multiple comparisons test were used for statistical analysis of the data (Prism 7 or 8 software). Significance threshold was set to be p<0.05. Data are presented in box plot with distribution of individual n’s and shown as mean ± standard deviation.

## Acknowledgements

We are grateful for Greif lab members and Gomperts lab members (Tammy Rickabaugh, Andrew Lund, Woosuk Choi, Chandani Sen) for their input on the experimental results and plans and the manuscript. We thank Kevin Boyen for technical experimental help. We also thank I. Weissman, D. Metzger and P. Chambon for mouse strains. Funding was provided by the Department of Army (W81XWH-18-1-0629 to D.M.G.), National Institute of Health (R35HL150766, R01HL125815, R01HL133016, R21NS088854 to D.M.G. and F32HL132532 to R.R.C.), American Heart Association (Established Investigator Award, 19EIA34660321 to D.M.G.).

## Author contributions

R.R.C. and D.M.G. conceived of and designed experiments, and R.R.C., I.K. and E.G. performed them. T.A. conducted scRNA-seq analysis, and N.K. provided infrastructure for this analysis. R.R.C and D.M.G. analyzed the results, and R.R.C, T.A. and D.M.G. prepared the figures. R.R.C, B.G. and D.M.G. wrote the manuscript.

