## Supplemental figs. and legends for "Dedifferentiated early postnatal lung myofibroblasts redifferentiate in adult disease"

**List of Supplementary Items:**

- Supplementary Figures 1-6
- Supplementary Legends

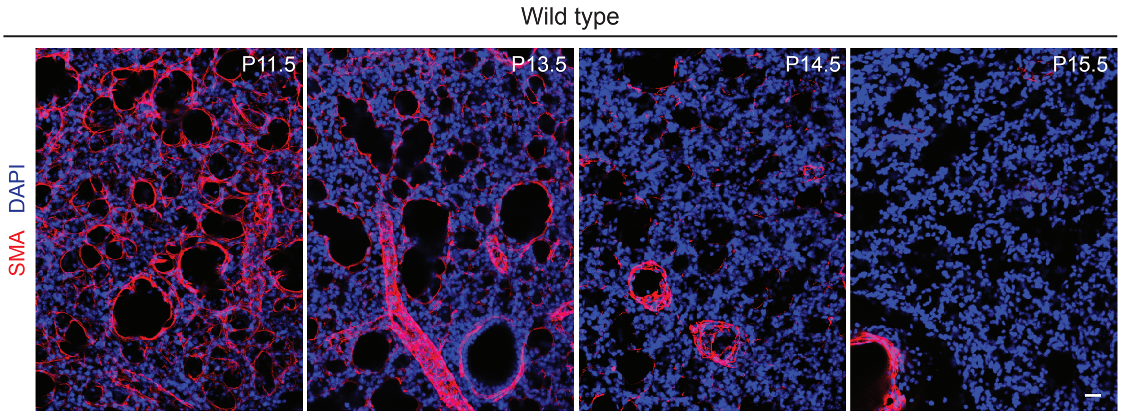
Wild type
SMA DAPI
P11.5
P13.5
P14.5
P15.5

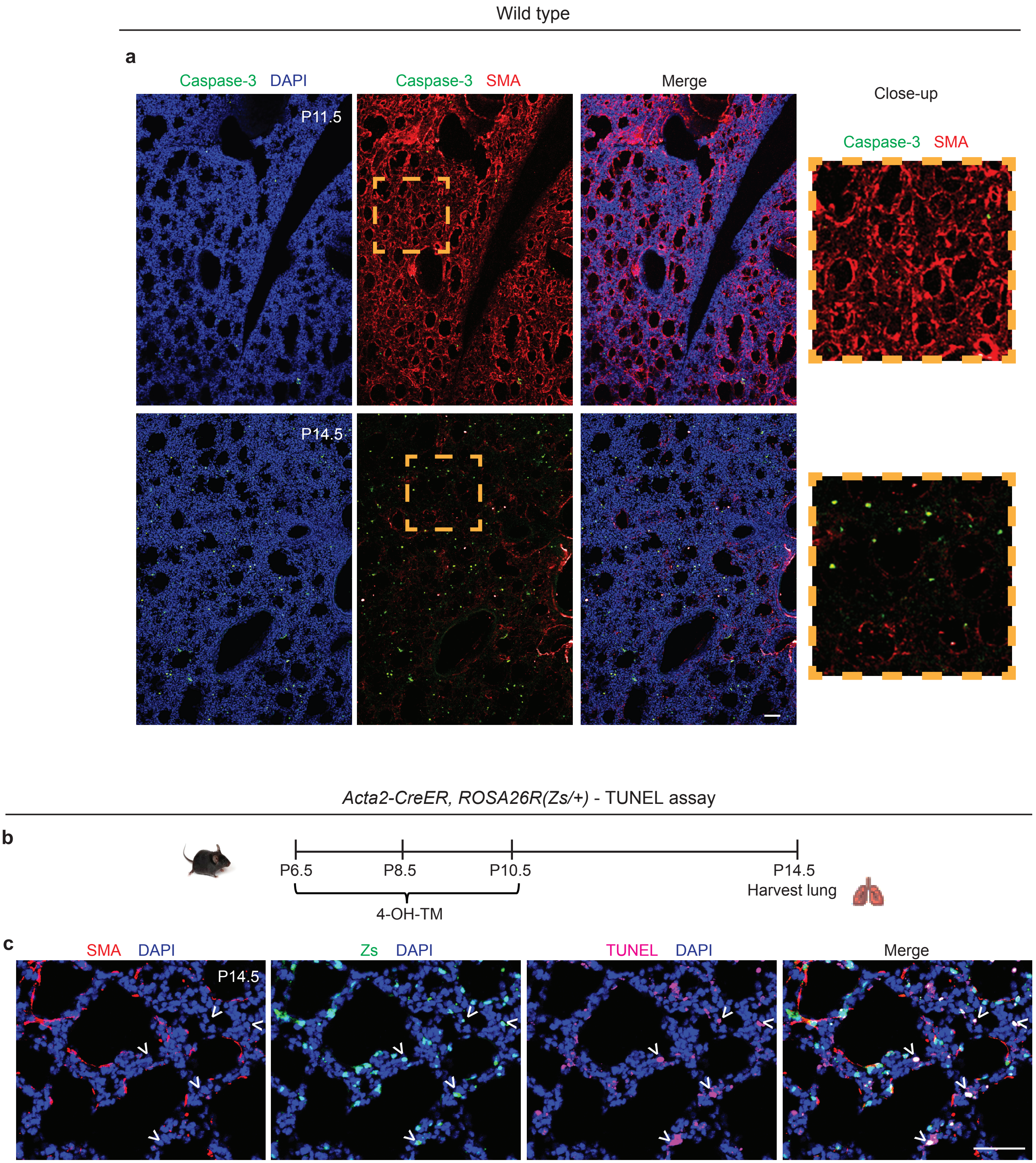
Wild type
a
Caspase-3 DAPI
Caspase-3 SMA
Merge
Close-up
Caspase-3 SMA
P11.5
P14.5
Acta2-CreER, ROSA26R(Zs/+) - TUNEL assay
b
P6.5
P8.5
P10.5
P14.5
Harvest lung
4-OH-TM
c
SMA DAPI
Zs DAPI
TUNEL DAPI
Merge
P14.5

*Acta2-CreER, ROSA26R(Zs/+)* - 4-OH TM, P6.5 to P10.5; analysis, P17.5

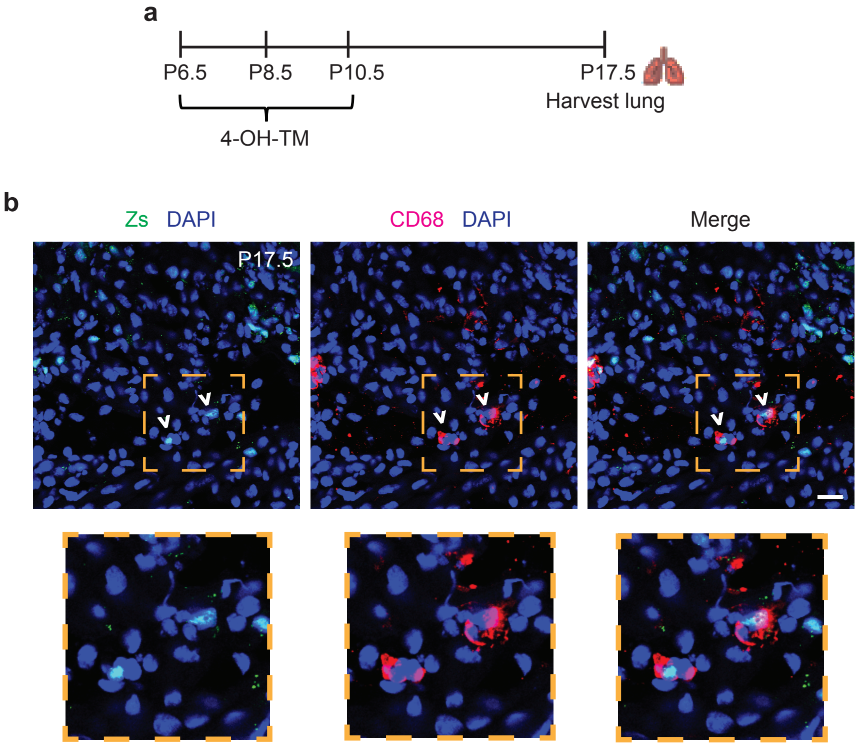

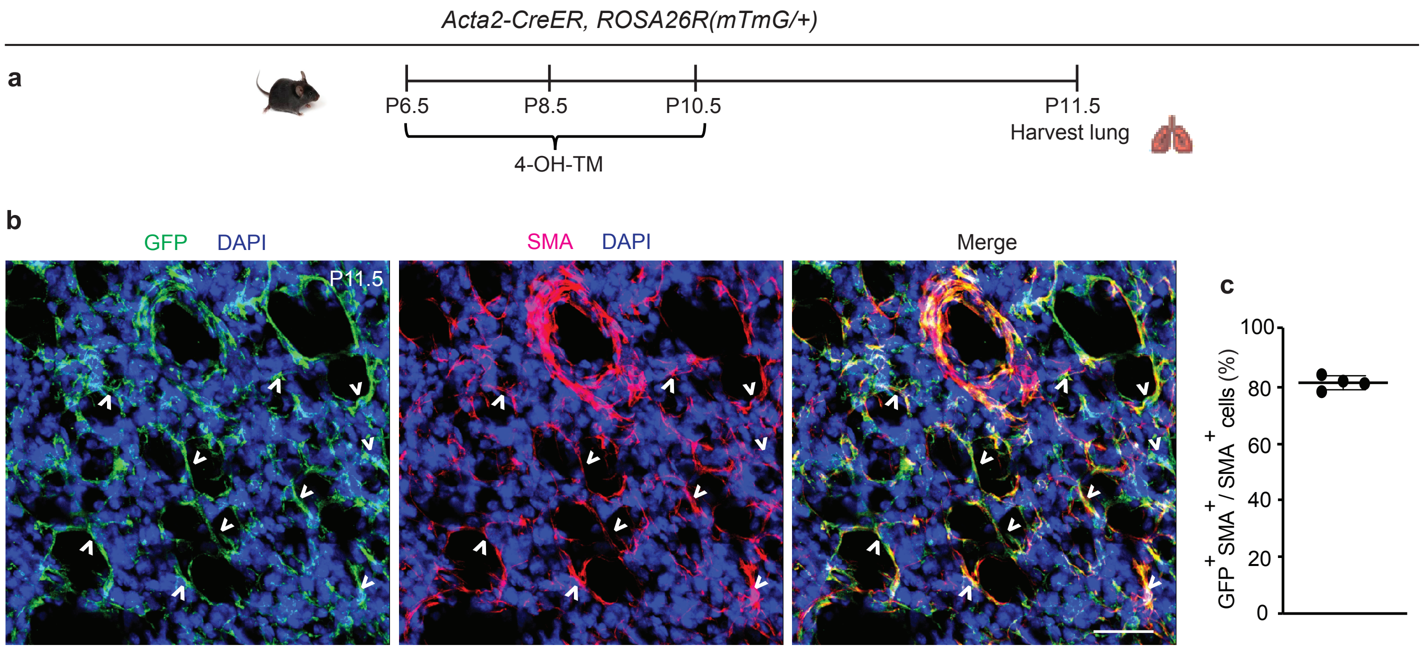
Acta2-CreER, ROSA26R(mTmG/+)
a
P6.5
P8.5
P10.5
P11.5
Harvest lung
4-OH-TM
b
GFP
DAPI
SMA
DAPI
Merge
P11.5
c
100
80
60
40
20
0
GFP+ SMA+ / SMA+ cells (%)

*Acta2-CreER, ROSA26R(mTmG/+)* - 4-OH TM, P6.5 to P10.5; analysis, P11.5

**a**

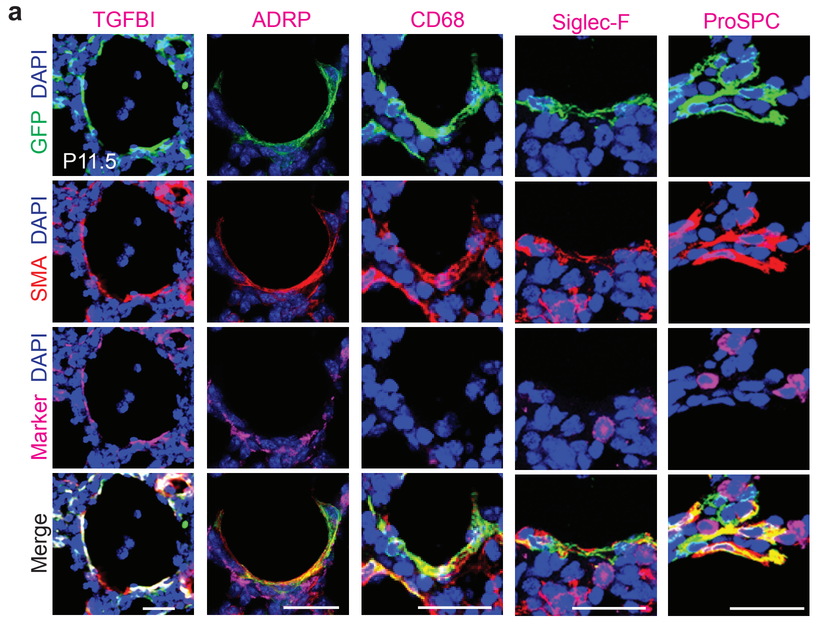


*Acta2-CreER, ROSA26R(mTmG/+)* - 4-OH TM, P6.5 to P10.5; analysis, P60.5

**b**

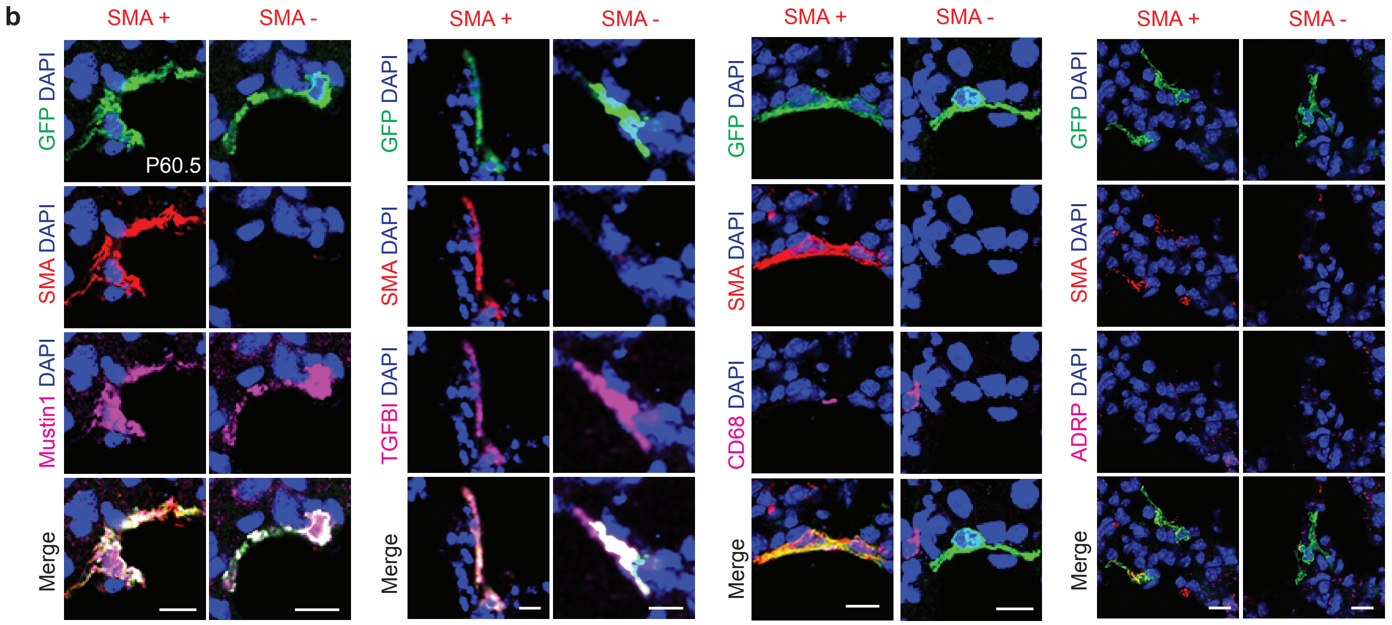

***Acta2-CreER, ROSA26R(mTmG/+)*** - 4-OH TM, P6.5 to P10.5; analysis, P60.5

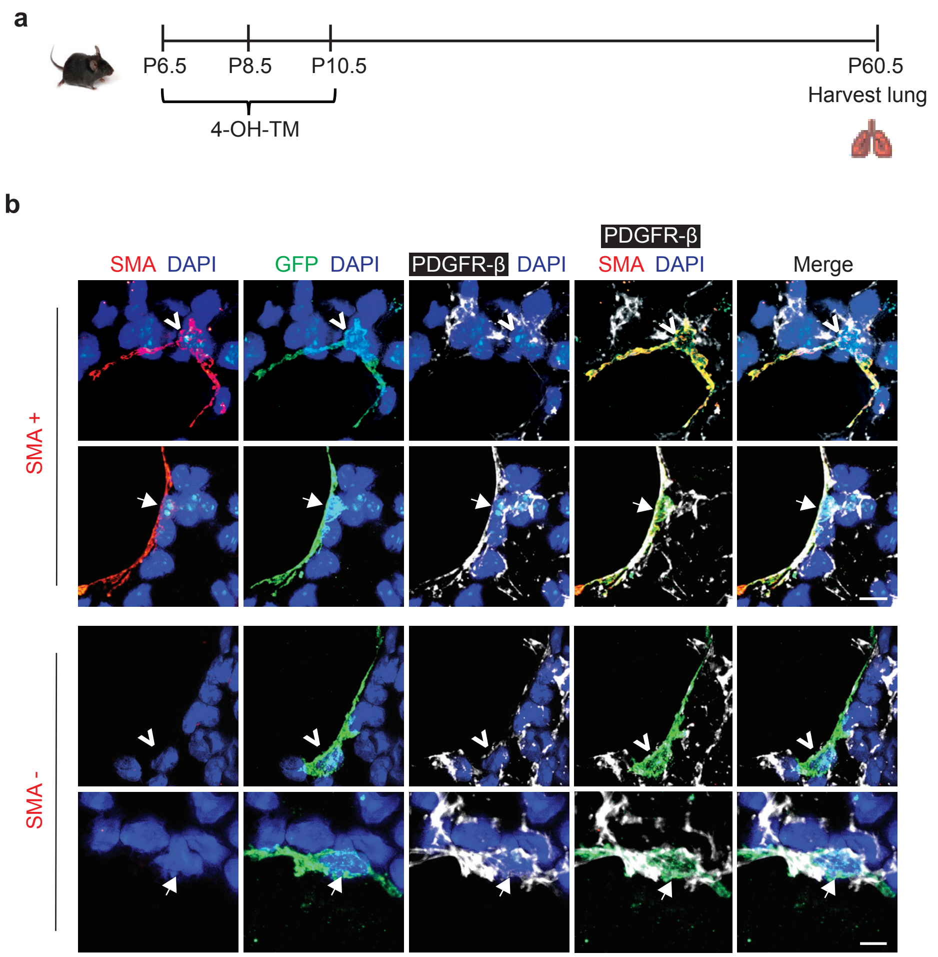

**Supplementary Figure Legends:**

**Figure S1. Time course of loss of SMA$^+$ myofibroblasts in the lung during postnatal alveolarization.** Lungs from wild type mice at indicated postnatal ages were harvested, cryosectioned and stained for SMA and nuclei (DAPI). Representative images are shown; n=5 mice per time point. Scale bar, 25 μm.

**Figure S2. SMA$^+$ myofibroblasts downregulate SMA before undergoing apoptosis. a,** Cryosections of the lungs of wild type mice at P11.5 and P14.5 stained for caspase 3 and nuclei (DAPI) are shown. Close-ups of boxed regions are displayed on the right; n=3 mice. **b, c,** *Acta2-CreER$^{T2}$, ROSA26R$^{(Zs/+)}$* mice were induced with 4-OH TM at P6.5, P8.5 and P10.5 and then euthanized at P14.5. Lungs were harvested and cryosectioned. In **b**, schematic of the experiment is shown. In **c**, cryosections were stained for SMA, TUNEL and nuclei (DAPI) and directly imaged for Zs (fate marker); open arrowheads indicate Zs$^+$TUNEL$^+$ cells. n=3 mice. Scale bars, 100 μm **(a)**, 50 μm **(c)**.

**Figure S3. Association of apoptotic myofibroblasts or their cell debris with macrophages.** *Acta2-CreER, ROSA26R$^{(Zs/+)}$* mice induced with 4-OH TM at P6.5, P8.5 and P10.5 were euthanized at P17.5. **a**, Schematic showing the experimental set-up. **b**, Lung cryosections were stained for CD68 (macrophage marker), Zs and nuclei (DAPI). Representative images are shown with close-ups in the boxed region below; open arrowheads indicate co-localization of Zs within CD68$^+$ cells. n=3 mice. Scale bar, 50 μm.

**Figure S4. Cell marking efficiency in *Acta2-CreER$^{T2}$*.** *Acta2-CreER$^{T2}$, ROSA26R$^{(mTmG/+)}$* mice were induced with 4-OH TM at P6.5, P8.5 and P10.5. Mice were euthanized at P11.5, and lungs were harvested. **a,** Schematic of the labeling and lung collection timeline is shown. **b**, Lung cryosections were stained for GFP (fate marker), nuclei (DAPI) and SMA, and imaged; open arrowheads indicate overlap of GFP and SMA. Scale bar, 50 µm. **c**, Labeling efficiency (% of $SMA^+$ cells that are $GFP^+$) was quantified from n=4 mice.

**Figure S5. Marker analysis at P11.5 and P60.5 of fate mapped early postnatal $SMA^+$ cells.** *Acta2-CreER$^{T2}$*, *ROSA26R$^{(mTmG/+)}$* mice were induced with 4-OH TM at P6.5, P8.5 and P10.5, and lungs were harvested at P11.5 or P60.5. Lung sections were stained for SMA, GFP (fate marker), nuclei (DAPI) and indicated markers (magenta). **a**, Representative images of $SMA^+GFP^+$ cells at P11.5 are shown; n=3-6. **b**, Lineage-traced $SMA^+$ and $SMA^-$ parenchymal cells at P60.5 are shown; n=4-6. Scale bars, 25 µm **(a)**, 10 µm **(b)**.

**Figure S6. PDGFR-β expression analysis of fate mapped early postnatal myofibroblasts at P60.5.** *Acta2-CreER$^{T2}$*, *ROSA26R*$^{(mTmG/+)}$ mice were induced with 4-OH TM at P6.5, P8.5 and P10.5, and lungs were harvested at P60.5. **a,** Schematic of experimental set up is shown. **b,** Vibratome sections of lungs at P60.5 were stained for SMA, GFP (fate marker), nuclei (DAPI) and PDGFR-β. Representative images of $SMA^+$ and $SMA^-$ cells are shown. Open arrowheads indicate PDGFR-β low cells, and arrows indicate PDGFR-β high cells; n=3 mice. Scale bars, 10 µm.
